# Optimal operation of parallel mini-bioreactors in bioprocess development using multi-stage MPC

**DOI:** 10.1101/2021.12.17.472671

**Authors:** Niels Krausch, Jong Woo Kim, Sergio Lucia, Sebastian Groß, Tilman Barz, Peter Neubauer, Mariano N. Cruz Bournazou

**Author notes:** Correspondence: Mariano N. Cruz Bournazou.

## Abstract

Bioprocess development is commonly characterized by long development times, especially in the early screening phase. After promising candidates have been pre-selected in screening campaigns, an optimal operating strategy has to be found and verified under conditions similar to production. Cultivating cells with pulse-based feeding and thus exposing them to oscillating feast and famine phases has shown to be a powerful approach to study microorganisms closer to industrial bioreactor conditions. In view of the large number of strains and the process conditions to be tested, high-throughput cultivation systems provide an essential tool to sample the large design space in short time. We have recently presented a comprehensive platform, consisting of two liquid handling stations coupled with a model-based experimental design and operation framework to increase the efficiency in High Throughput bioprocess development. Using calibrated macro-kinetic growth models, the platform has been successfully used for the development of scale-down fed-batch cultivations in parallel mini-bioreactor systems. However, it has also been shown that parametric uncertainties in the models can significantly affect the prediction accuracy and thus the reliability of optimized cultivation strategies. To tackle this issue, we implemented a multi-stage Model Predictive Control (MPC) strategy to fulfill the experimental objectives under tight constraints despite the uncertainty in the parameters and the measurements. Dealing with uncertainties in the parameters is of major importance, since constraint violation would easily occur otherwise, which in turn could have adverse effects on the quality of the heterologous protein produced. Multi-stage approaches build up scenario tree, based on the uncertainty that can be encountered and computing optimal inputs that satisfy the constrains despite of such uncertainties. Using the feedback information gained through the evolution along the tree, the control approach is significantly more robust than standard MPC approaches without being overly conservative. We show in this study that the application of multi-stage MPC can increase the number of successful experiments, by applying this methodology to a mini-bioreactor cultivation operated in parallel.

## 1. Introduction

### 1.1 High-throughput bioprocess development

The development of a process in biotechnology is enormously time-consuming. Not only does it take a long time to find a suitable strain from a huge collection of strains, but the identification of optimal process conditions for this strain also takes a lot of time and resources. Especially due to the increased demand for resource-saving and environmentally friendly bioproducts, the requirements for faster process development are also increasing. Miniaturization and parallelization are two recent technologies that can accelerate throughput in screening processes and process development (Hemmerich et al., 2018). In particular, the use of liquid handling stations and the application of computer-aided and model-based methods could make a decisive contribution to holistic and faster process development (Hans et al., 2020). These methods have already been used for optimal experimental re-design (Cruz Bournazou et al., 2017, Barz et al., 2018) as well as process insights and faster strain phenotyping (Anane et al., 2019).

### 1.2. Advanced control methods of bioprocesses

Fed-batch is still the most widely used process strategy in bioprocessing. Here, the change of the feeding rate plays an important role. Classically, feeding sequences after the batch phase follow an exponential course, based on a previously defined growth rate. However, it can easily happen that the selected growth rate is either too low and thus the process takes a long time to reach a certain cell density or that the feeding rate was selected too high and oxygen limitation can occur. This is a problem for various processes, as it causes unwanted stress reactions and can significantly influence the yield (Baez & Shiloach, 2014). Model predictive control is a method that has been widely used in the chemical industry and has also shown good success in several simulation studies in bioprocess engineering but has had little application in real experiments (Rathore et al., 2021). This is mainly due to the fact that good estimates for the underlying parameters must be available for the model. Finding good values for this is often time consuming and the values are subject to large uncertainties. We show here our implementation of a combination of moving horizon estimation for parameter estimation and multi-stage model predictive control. This strategy provides rough estimates for the parameters for new strains where no information is available in a short time, and multi-stage MPC is then used to compute an optimal feeding regime while explicitly taking into account the uncertainty of such parameters. As more information, in the form of measurements, is gathered, the parameter estimation can be improved and the multi-stage MPC can include this information online to improve the performance while ensuring robust process control and avoiding oxygen limitation. Since the dissolved oxygen tension (DOT) is measured with a first order delay, it is not possible to control the process optimally by means of a PID controller, since the violation of the constraint always occurs after a glucose pulse has been added. In this respect, a model-based control with prediction of whether a selected pulse satisfies the constraint is necessary. This is another reason to take the uncertainties of the parameter values into account, as this approach has proven to be significantly more robust than previous approaches (Lucia et al., 2013). By using our fully automated liquid handling station, we can quickly take many measurements to adapt our model well to the new strain and at the same time use the outputs of the MPC to optimally adjust the process online. The multi-stage approach in particular makes it possible to still ensure robust process control even with inaccurate estimated values for the parameters.

## 2. Materials and Methods

Initial testing experiments for the parameter estimation were conducted on our high-throughput bioprocess development platform, although the actual multi-stage experiments were performed in silico. The platform comprises two liquid handling stations (Freedom Evo 200, Tecan, Switzerland; Microlab Star, Hamilton, Switzerland), a mini bioreactor system (48 BioReactor, 2mag AG, Munich, Germany) and a Synergy MX microwell plate reader (BioTek Instruments GmbH, Bad Friedrichshall, Germany). With this setup, the platform is capable to perform up to 48 cultivations in parallel. DOT and pH are measured online, while the concentrations of the other state variables, i.e. glucose, acetate and biomass were measured at-line using enzymatic kits. The reader is referred to (Haby et al., 2019) for a detailed description of the facility and sampling procedure. The multi-rate sampling frequency of the different states poses another challenge for the optimization, as DOT is measured every 60 s, while the other state variables are only measured every 20 min.

The MHE part for parameter and initial state estimation can be generally formulated according to (1), subject to (2).

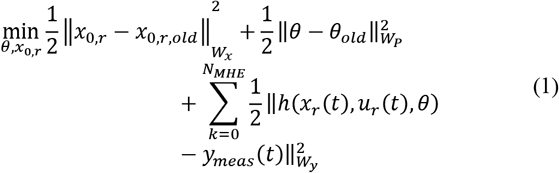

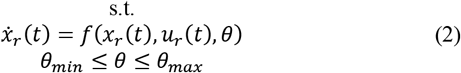

The estimate for the states at the initial point of the window are then denoted byx_0,r,old_, the current parameter vector *θ*, the parameter vector from the previous horizon *θ_old_*. The squared norm is applied to all calculations and the optimizer tries to minimize the deviations over the time window of length *N_MHE_* of the measurements *y_meas_* and the predicted outputs *h*(*x_r_*(*t*), *u_r_*(*t*),*θ*), considering the inputs *u_r_*(*t*) and current parameter vector. Each norm is further weighted by the factors *W_x_*, *W_p_*, *W_y_*.

The optimization problem that is solved to compute the optimal inputs via MPC is:

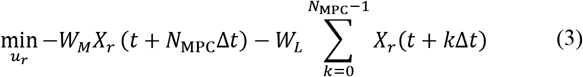

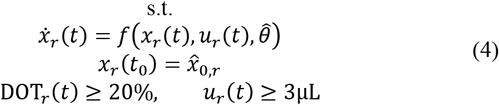

Where *W_V_* and *W_L_* denote the weightings for the terminal- and the stage-cost, respectively. The goal was to maximize the biomass *X_r_* in the shortest time while at the same time avoiding oxygen limitation, i.e. having a level of at least 20% dissolved oxygen in the medium.

The problem was solved using the do-mpc software which utilizes orthogonal collocation on finite elements to discretize the system, which can then be solved using NLP optimizers (Lucia et al., 2017). One main challenge when operating this system is the non-continuous property of the inputs. Since glucose feeding is bolus-like, concentrations and volumes are changed at discrete time points. To deal with this, mass balances must be solved for the respective pulse additions, and it is mathematically more difficult to solve the system. The reader is referred to (Kim et al., 2021) for an in-depth description of the underlying model and description of the optimization process. To consider the uncertainties in the parameters, a scenario tree is built. Here, each branch of the tree represents a possible combination of the uncertain parameter combinations. At each node, the optimization problem is solved again considering the current parameter uncertainty and an optimal trajectory is identified. When the parameter uncertainty is large, this approach yields much better constraint satisfaction than nominal MPC approaches. In this contribution, we tested the multi-stage approach to consider the uncertainties for a prediction horizon of 180 min and a robust control horizon of 10 min. A schematic description of the scenario tree with arbitrary lengths is depicted in Figure 1.

**Figure 1:**
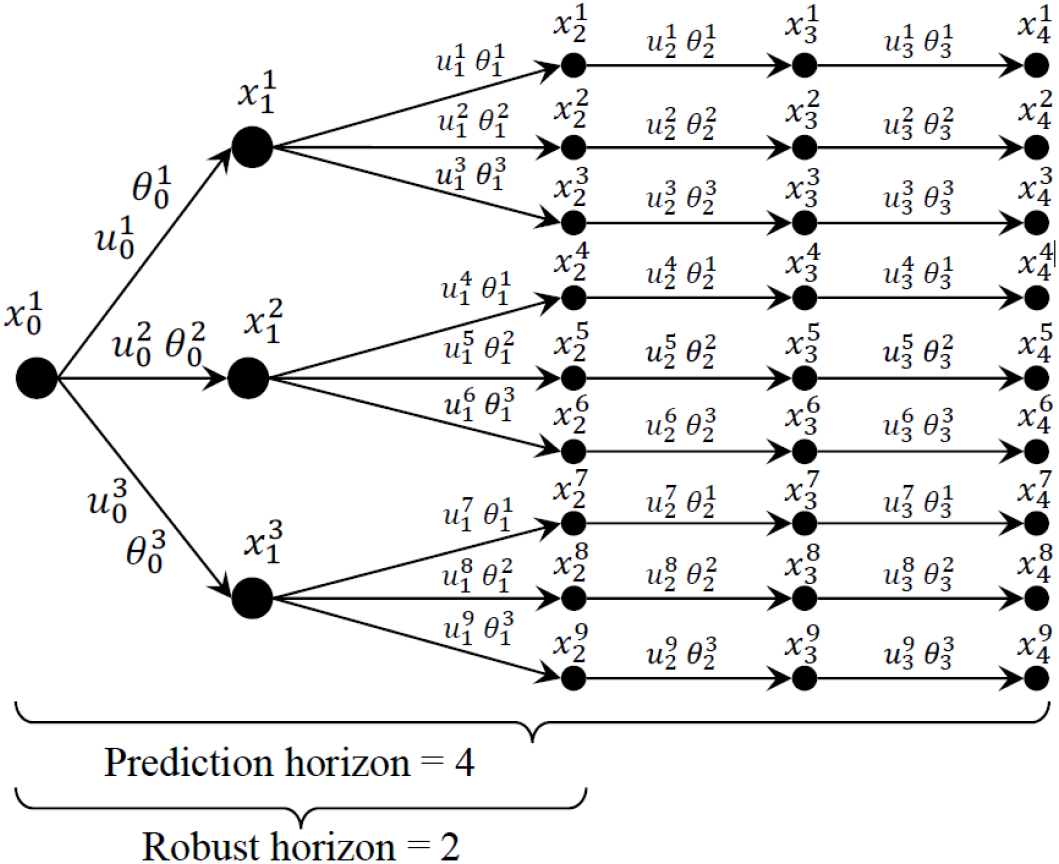
Scenario tree. Schematic overview about how the tree is built with an arbitrary prediction horizon of 4 and a robust horizon of 2. Uncertainties are considered until the second node.

## 3. Results and Discussion

Based on our previous work, where we showed the advantage of using MPC to obtain higher biomass while avoiding oxygen limitation compared to a predefined feeding regime (Krausch et al., 2020), we extended our work to account for the uncertainties in the parameters. Compared to the nominal MPC, multi-stage MPC offers the opportunity to consider the inherent uncertainties of the model parameters to be more robust. This is especially important to ensure that the defined constraints are not violated. Especially for processes with a large uncertainty in the parameters or difficult to predict dynamics, this offers a much better possibility of process control. In this setup, the framework was used to consider the uncertainties of three important model parameters, namely the maximum substrate uptake rate *q_S,max_*, the yield coefficient of biomass per substrate excluding maintenance *Y_XS,em_* and the volumetric oxygen transfer coefficient *k_L_a*. All these parameters can affect the oxygen consumption and hence easily lead to constraint violation. Choosing the right parameters in the scenario tree could be supported by using a subset selection method with sensitivity analysis or further data-driven approaches like PCA (Thombre et al., 2019). Many studies have shown the advantage of advanced control methods such as MPC, but have mainly been studying the nominal case for optimization (Chang et al., 2016; Ulonska et al., 2018). As depicted in Figure 2, the multi-stage MPC calculates a lower feeding rate compared to the nominal MPC, since it considers the uncertainties in the parameters. The nominal MPC can actually only ensure that the constraints are not violated if the parameters are well estimated and have very low uncertainty. The multi-stage approach, on the other hand, can identify an optimal feeding regime and account for the strength of uncertainties via a feedback loop. This is particularly advantageous for processes about which little information and measurement data are available like in early screening of new strains and therefore the parameter uncertainties are correspondingly large. Accordingly, taking these uncertainties into account leads to a lower feeding behavior in contrast to the nominal MPC, but to a much more robust process behavior. In particular, the system can react quickly in an unpredictable manner if the constraint is violated, since oxygen limitation also has a strong influence on the expression of numerous genes and thus could strongly change the dynamics of the system.

**Figure 2:**
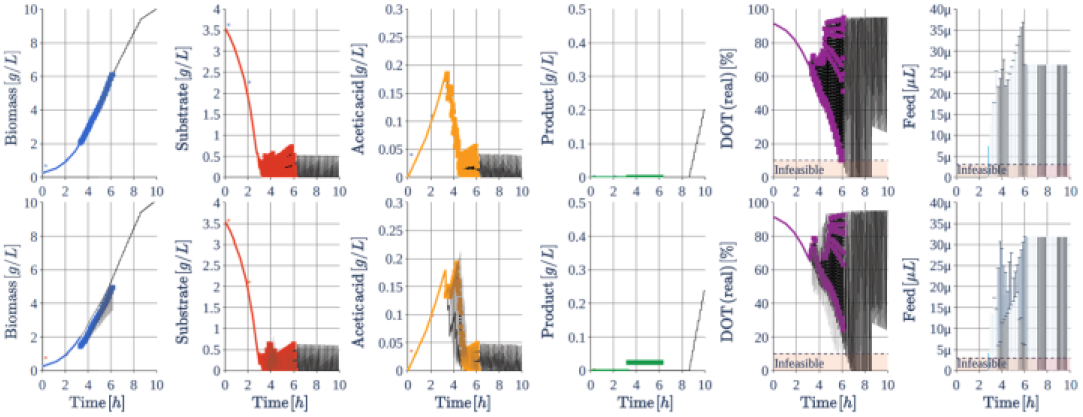
Comparison of nominal (upper) and multi-stage (lower) MPC at 3 h. The multi-stage approach guarantees constraint satisfaction even when the uncertainties are considered. Uncertainties for the DOT and feed are shown as shaded plots and error bars, respectively.

## 4. Conclusion and outlook

In this study, we have shown that a robust multi-stage MPC approach that takes into account the uncertainties in the parameters is even better than a nominal MPC. Violation of the constraint is a problem for many bioprocesses and there are various approaches to control bioprocesses. With the predictive approach, the constraints can be met well before they are violated, and the process can be guided to its optimum, so constraints are satisfied even in case of high uncertain parameters. However, the computational effort increases significantly with the number of parameters and the underlying uncertainties, so careful consideration must be given to which parameters the uncertainties should be considered for. Future work will need to deal with this problem by being more flexible in terms of what parameter uncertainties are considered and how large the prediction horizon should be, since the problem size is growing exponentially with a larger robust horizon or number of uncertain parameters to account for. Another aspect is to represent the product formation with the help of data-driven models, so that the process can also be optimized in this respect.

## Acknowledgments

We thank Felix Fiedler for the development of the do-mpc software. We gratefully acknowledge the financial support of the German Federal Ministry of Education and Research (BMBF) (project no. 01QE1957C – BioProBot and 01DD20002A – 568 KIWI Biolab).

## Notes

### Competing Interest Statement

The authors have declared no competing interest.

